# TMC5 promotes lung adenocarcinoma cell proliferation, migration and invasion by regulating epithelial-mesenchymal transition

**DOI:** 10.64898/2025.12.23.696250

**Authors:** Danxiong Sun, Rufang Li, Yunhui Zhang

**Author notes:** CORRESPONDENCE: Yunhui Zhang, emai.

## Abstract

**Background/purpose:** In recent years, evidence has indicated that transmembrane channel-like 5 (TMC5) plays an important role in various cancers. Previously, one study showed that TMC5 was highly expressed in lung adenocarcinoma (LUAD), but its mechanism in LUAD remained unclear. In this study, we aimed to investigate the effects of TMC5 and its underlying mechanisms in LUAD progression.

**Methods:** The expression level of TMC5 in LUAD tissues was detected by real-time quantitative PCR. TMC5 was knocked down in A549 and H1437 cell lines using lentiviral transfection. The oncogenic effect of TMC5 on LUAD cell proliferation was evaluated by CCK-8 assay and colony formation assay. Transwell assay was employed to assess migration and invasion.

**Results:** TMC5 was overexpressed in LUAD tissues compared with normal tissues. Suppression of TMC5 expression reduced proliferation, migration, and invasion, while also decreasing the expression of mesenchymal markers and increasing epithelial markers in A549 and H1437 cells. Conversely, higher TMC5 expression significantly increased the expression of mesenchymal markers and decreased epithelial markers.

**Conclusion:** TMC5 facilitates the malignant behaviors of LUAD cells by regulating epithelial-mesenchymal transition, suggesting its potential as a promising diagnostic and therapeutic target.

## Introduction

The leading cause of cancer death worldwide is lung cancer (1). The most common pathological type of lung cancer is adenocarcinoma. In recent years, significant breakthroughs have been made in targeted molecular therapy and immunotherapy for lung adenocarcinoma (LUAD). However, the five-year survival rate for LUAD remains unsatisfactory. Metastasis is not only the leading cause of death in LUAD, but also a major hurdle for its treatment. Therefore, further elucidation of the molecular mechanism of LUAD metastasis, searching for new metastasis-related factors have important clinical significance.

Transmembrane channel-like (TMC) proteins are a gene family of evolutionarily conserved ion channel-like membrane proteins. It was reported that the TMC gene plays an important role in various diseases, including hearing loss, cancer, and fatty liver disease (2). The functional research of TMC5 in human cancers is largely unexplored. In recent years, evidence has indicated that TMC5 plays a role in prostate cancer, hepatocellular carcinoma,and gastric cancer. Through cell cycle regulation, TMC5 promotes prostate cancer cell proliferation (3). The study by Li et al. demonstrated that TMC5 promotes hepatocellular carcinoma progression by regulating epithelial-mesenchymal transition (EMT) (4). Functional assays conducted by Wei et al. demonstrated that TMC5 promoted gastric cancer cell proliferation, migration, and invasion in vitro, as well as tumor growth and metastasis in vivo (5).

Previously, one study of TMC5 showed that it was highly expressed in LUAD (it was non-significant in small cell lung cancer and squamous cell carcinoma of the lung), but the mechanism of TMC5 in LUAD was still unclear (6). In the present study, we first assessed the biological function of TMC5 in LUAD, and explored its mechanism of action. Our results highlight the underlying mechanisms of TMC5 regulation in LUAD.

## Materials and Methods

### Sample collection

The samples for this study were derived from archived specimens. The data were accessed for research purposes from December 1, 2024, to December 31, 2024. All data had been fully anonymized prior to access. The authors had no access to information that could identify individual participants during or after data collection.

To detect the expression of TMC5 in LUAD tissue, the leftover RNA from the samples that had undergone transcriptome sequencing (n=16) was used for reverse transcription and qPCR.

Our study was approved by the Ethics Committee of the First People’s Hospital of Yunnan Province (Authorization number: KHLL2024-KY282). The informed consent was obtained from all subjects, and all patients have signed the informed consent forms.

### Cell culture

A549 and NCI-H1437 cells, all derived from human LUAD, were purchased from cell bank of Chinese Academy of Science, Shanghai, in China. A549 cells were cultured in F12K medium (Servicebio, Wuhan, China), and NCI-H1437 cells were cultured in RPMI 1640 medium (servicebio, Wuhan, China). All of the media contain 10% fetal bovine serum (Gibco), 1% penicillin-streptomycin. The cells were maintained at 37°C, in a humidified incubator with 5% CO2. Cells in the logarithmic growth phase were used for experiments.

### RNA extraction, reverse transcription, and qRT-PCR

Total RNA was extracted from the cells using a FreeZol Reagent Kit (Vazyme, Nanjing, China) according to the manufacturer’s instructions. Synthesized cDNA was prepared with a Veriti 96-well thermal cycler (Applied Biosystems, Thermo Fisher Scientific, USA) for qRT-PCR using the Lepgen-96 Real-Time PCR System (LEPU, Beijing, China) with ChamQ Universal SYBR qPCR Master Mix (Vazyme, Nanjing, China). The relative expression of *TMC5* was determined with the 2^-△△Ct^ method after normalization to the expression of β-actin.

The primers for *TMC5* were: forward, 5′-ACTCTGGTGTGGATTGGCATCTTC-3′ and reverse, 5′-GCTCGGAGGCTGGAAATTCATCAT-3′.

### Western blot

Total proteins were prepared by lysing the cells with RIPA buffer containing the protease inhibitor cocktail (Beyotime, Shanghai, China) on ice for 15 min. Then the cell lysates were centrifuged at 4°C, 12000 g for 5 min. Protein concentrations were determined using the BCA Protein Assay Kit (Beyotime, Shanghai, China) according to the manufacturer’s instructions.

All the protein specimens were split up using an SDS polyacrylamide gel (Vazyme, Nanjing, China) and then the target protein were transferred to the 0.22-μm polyvinylidene difluoride (PVDF) membrane (Millipore, USA). The PVDF membranes were blocked at room temperature with 5% skim milk for 1.5 h. The PVDF membrane was incubated with the corresponding primary antibody (Table S1). This was followed by incubation with secondary antibody (HUABIO, Hangzhou, China). The primary and secondary antibodies were all incubated at room temperature for 1h. Anti β-actin served as control.

The protein bands were visualized using ECL (Servicebio, Wuhan, China) and collected by the ChemiDocTm XRS Molecular Imager System (Bio-Rad, USA). The Image J software was used to analyze the band densities.

### Transfection assay

Short hairpin RNA constructs against TMC5 (shTMC5) were cloned in JN0038-pLUX-U6 vector using BsmB1 cloning sites. The target sequence for TMC5-specific shRNA was 5’-CCAGCTCACTTGTTCTGGAAA-3’. The overexpression lentiviral vector targeting TMC5 (pLV[Exp]-Puro-EF1A>TMC5) was designed and synthesized by VectorBuilder (Guangzhpu, China), and the empty plasmid was used as a negative control. Lentiviral particles were produced by co-transfection of HEK293T with the packaging plasmid psPAX2, the envelope plasmid pLP-VSV-G, and the respective vector. Virus was collected 48h after transfection.

Polybrene (Beyotime, Shanghai, China) was used for cell transfections according to the manufacturer’s instructions. The lentiviral-transfected cells were cultured in medium with puromycin for 3 days to select stable cell lines. The puromycin concentrations in the culture medium for A549 and NCI-H1437 cell lines were 1.5 μg/mL and 2.0 μg/mL, respectively.

### Cell Proliferation Assay

Cell Counting Kit-8 (CCK-8, MeilunBio, Dalian, China) was used to determine the viability of cells. Cells were plated in 96-well plates at 1,000 cells per well. Following 24, 48, 72, 96 and 120 h incubation, the medium was removed from the wells and 100 µl of medium containing 10% CCK-8 were added to each well and then incubated for 1 h at 37°C. The optical density (OD) at 450 nm was measured to estimate the cell proliferation.

### Colony formation assay

For the colony formation assay, cells were seeded in 6-well plates (100 cells/well) and cultured for 14 days. The resulting colonies were fixed with 4% formaldehyde for 30 min, and then stained with 1% crystal violet for 20 min. Finally, the number of clones was counted to evaluate cell proliferation.

### Transwell assay

Transwell chambers (Abbkine, Wuhan, China), with or without Matrigel (Corning, USA), were applied for examination of cell invasion and migration, respectively. Matrigel was prepared according to the manufacturer’s instructions. Transfected A549 or H1437 cells were suspended in serum-free medium and were placed onto the upper chambers of Transwell. Medium containing 10% FBS was plated into the lower chambers. Non-migratory or noninvasive cells were wiped out after 48 h. The culture chamber was fixed with 4% paraformaldehyde for 30 min, and stained with 1% crystal violet for 20min.

## Statistical analysis

Statistical analyses were performed using GraphPad Prism 8 (GraphPad Software, San Diego, CA, USA). All assays were repeated three times. The data are presented as mean ± standard deviation (SD) and analyzed by Student’s t-test or two-way ANOVA. Differences were considered statistically significant at p < 0.05.

## Results

### TMC5 is overexpressed in human LUAD

Analysis of The UALCAN database showed that the protein of TMC5 is highly expressed in LUAD (Fig. 1A). We performed reverse transcription and qPCR using the leftover RNA from the samples that had undergone transcriptome sequencing (n=16). The results demonstrated that TMC5 mRNA was overexpressed in LUAD tissues compared with normal tissues (Fig. 1B). These data suggested that TMC5 might play an important role in lung tumorigenesis.

**FIGURE 1.**
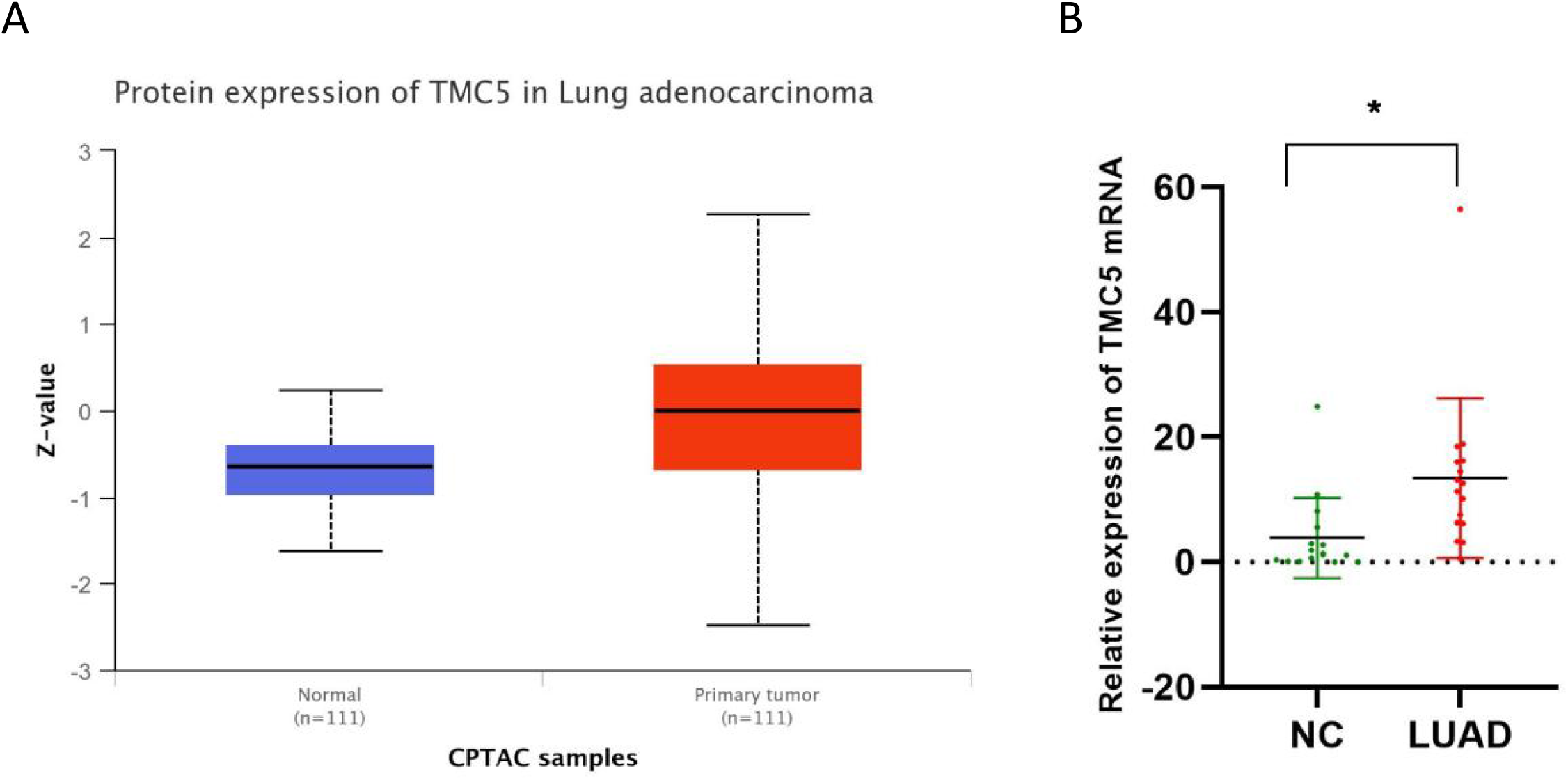
Expression of TMC5 in Lung Adenocarcinoma. (A) The protein of TMC5 is highly expressed in LUAD ( From the UALCAN database). (B) Compared with normal tissues, TMC5 mRNA was overexpressed in LUAD tissues. LUAD, Lung Adenocarcinoma. NC, normal control. *P < 0.05.

### Knockdown of TMC5 attenuated proliferation of LUAD cells

Stable TMC5-knockdown A549 and NCI-H1437 cell lines were successfully established via lentiviral transduction followed by puromycin selection (Fig. 2A,B). CCK-8 and clonogenic assays were performed to evaluate the effect of knocking down TMC5 expression on cell proliferation. The results show that TMC5 silencing decreased the absorbance reading at 450 nm, indicating that cell proliferation was attenuated (Fig. 2C-D). Similarly, colony formation was reduced in TMC5 knockdown cells (Fig. 2E-J), which indicated that blocking TMC5 expression dramatically reduces cell proliferation.

**FIGURE 2.**
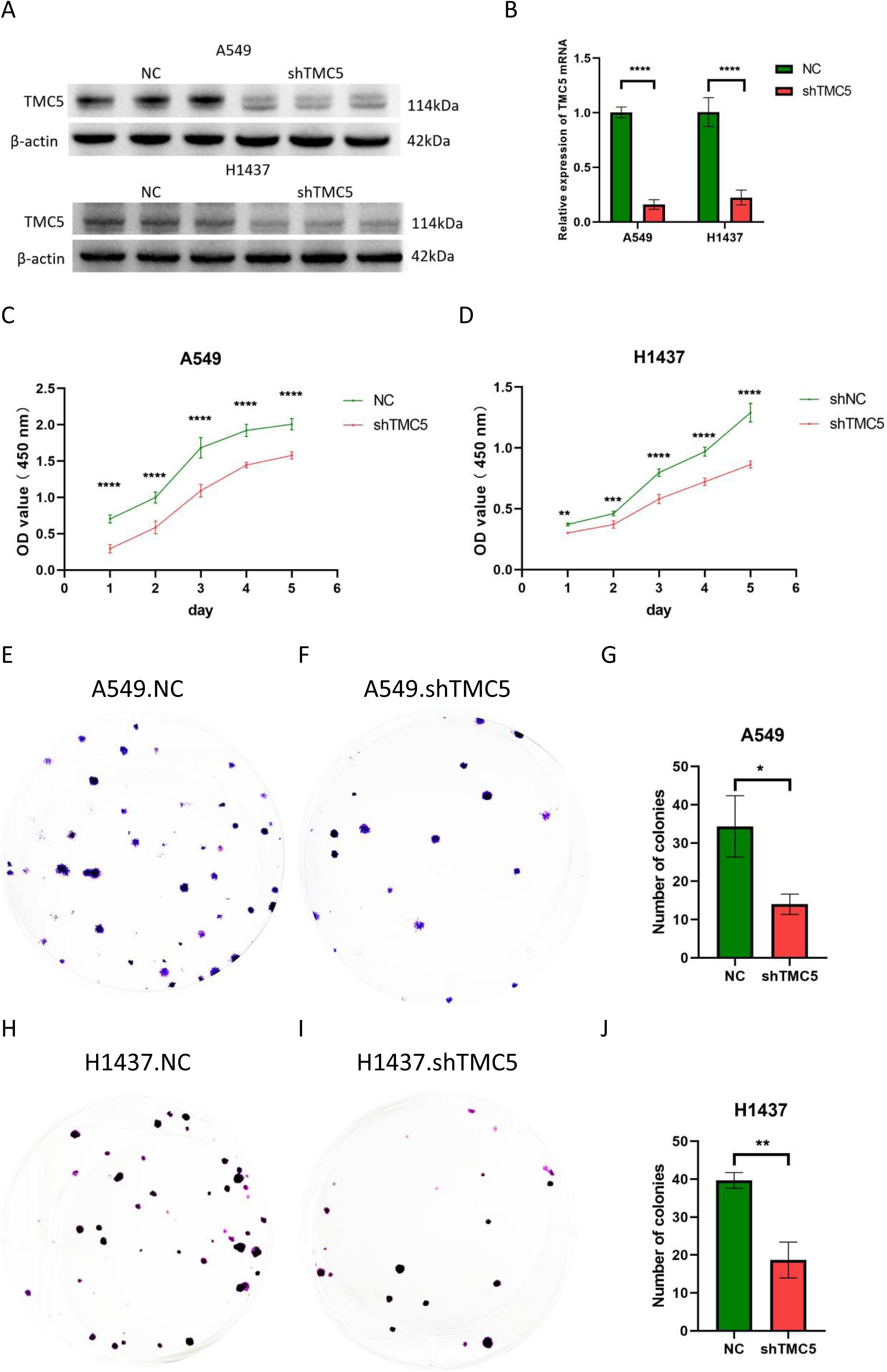
Effects of TMC5 Knockdown on the Proliferation of Lung Adenocarcinoma Cells. (A, B) qPCR and Western blot results showed that both mRNA and protein levels of TMC5 were downregulated in the knockdown cell lines. (C, D) CCK-8 assays demonstrated that the proliferation ability of A549 and NCI-H1437 cells was attenuated in TMC5 knockdown cells. (E-J) Colony formation assay results showed the impact of TMC5 knockdown in A549 and NCI-H1437 cells. NC: Normal control. shTMC5: Knockdown of TMC5 via short hairpin RNA ( shRNA). *P < 0.05, **P < 0.01, ***P < 0.0001, ****P < 0.00001.

### Knockdown of TMC5 inhibited LUAD cell migration and invasion

Transwell assay was used to evaluate the migration and invasion ability of LUAD cells. The number of migrated cells were decreased after TMC5 knockdown in A549 and NCI-H1437 cells (Fig. 3A-F). For invasion ability, the results were approximately similar to migration results (Fig. 4G-L). These results indicated that the function of TMC5 was to promote migration and invasion ability in A549 and NCI-H1437 cells.

**FIGURE 3.**
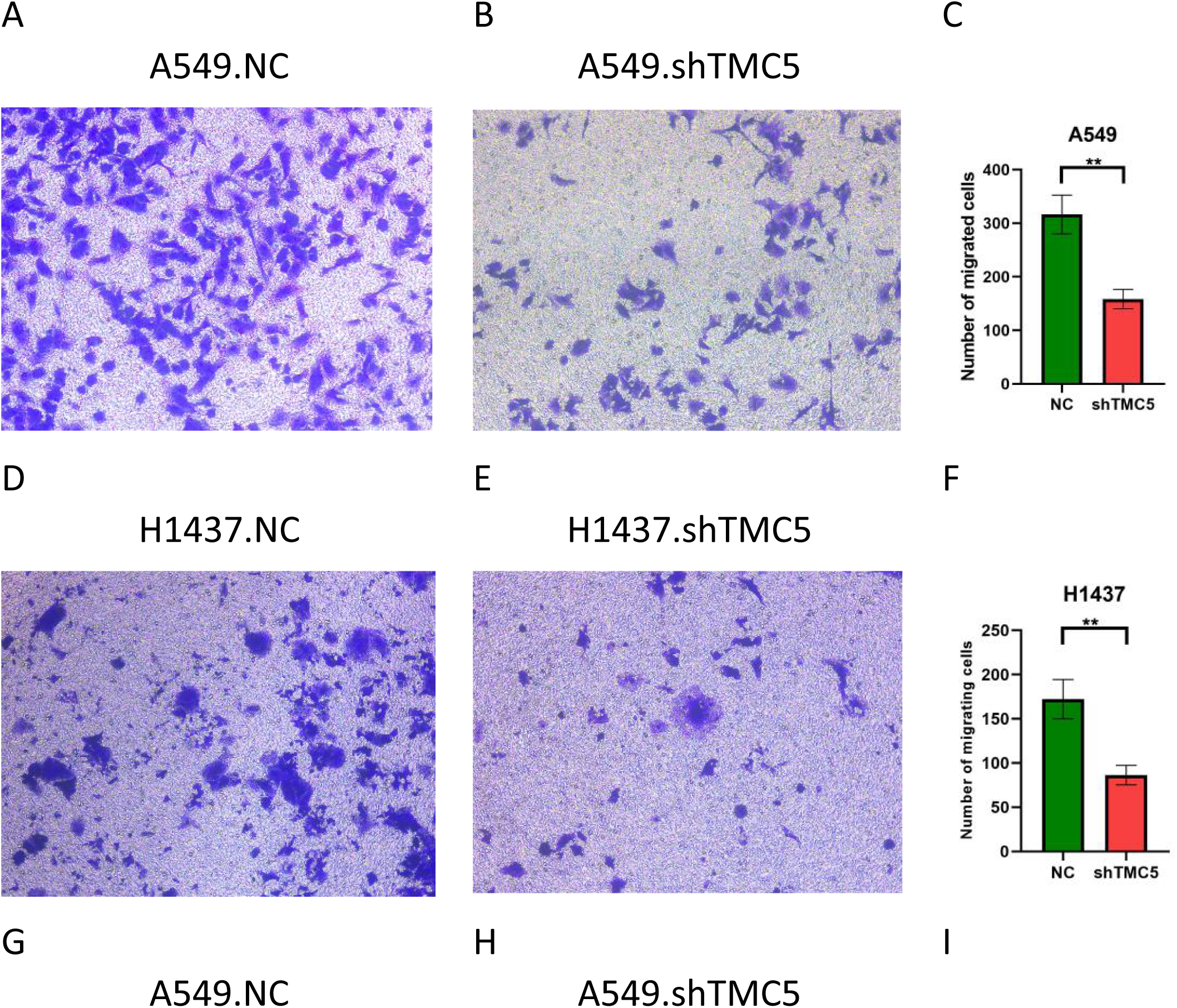
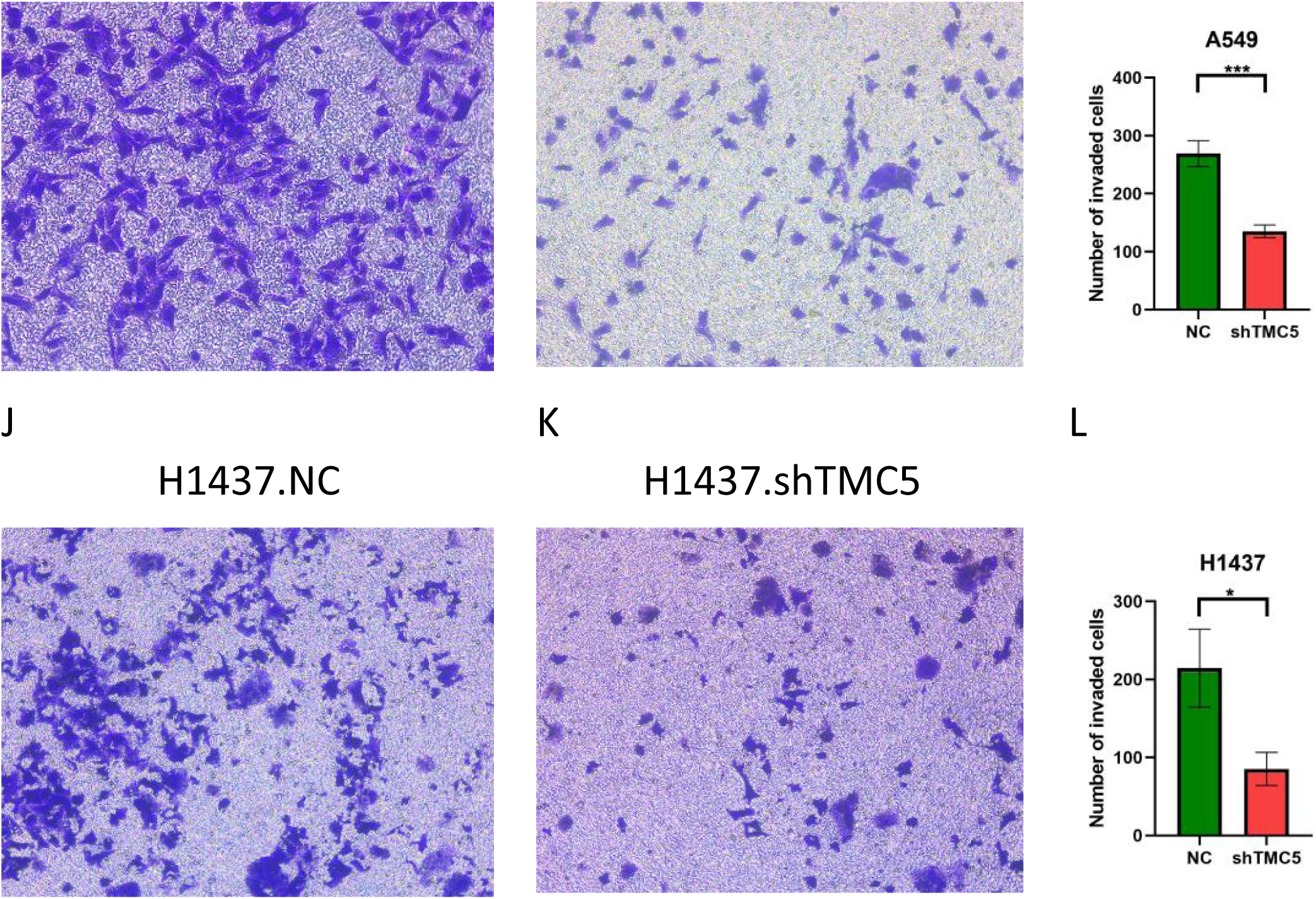
Effects of TMC5 Knockdown on the Migration and Invasion of Lung Adenocarcinoma Cells. (A-F) Representative images and results of Transwell migration assays. (G-L) Representative images and results of Transwell invasion assays. NC: Normal control. shTMC5: Knockdown of TMC5 via short hairpin RNA ( shRNA). *P < 0.05, **P < 0.01, ***P < 0.0001.

**FIGURE 4.**
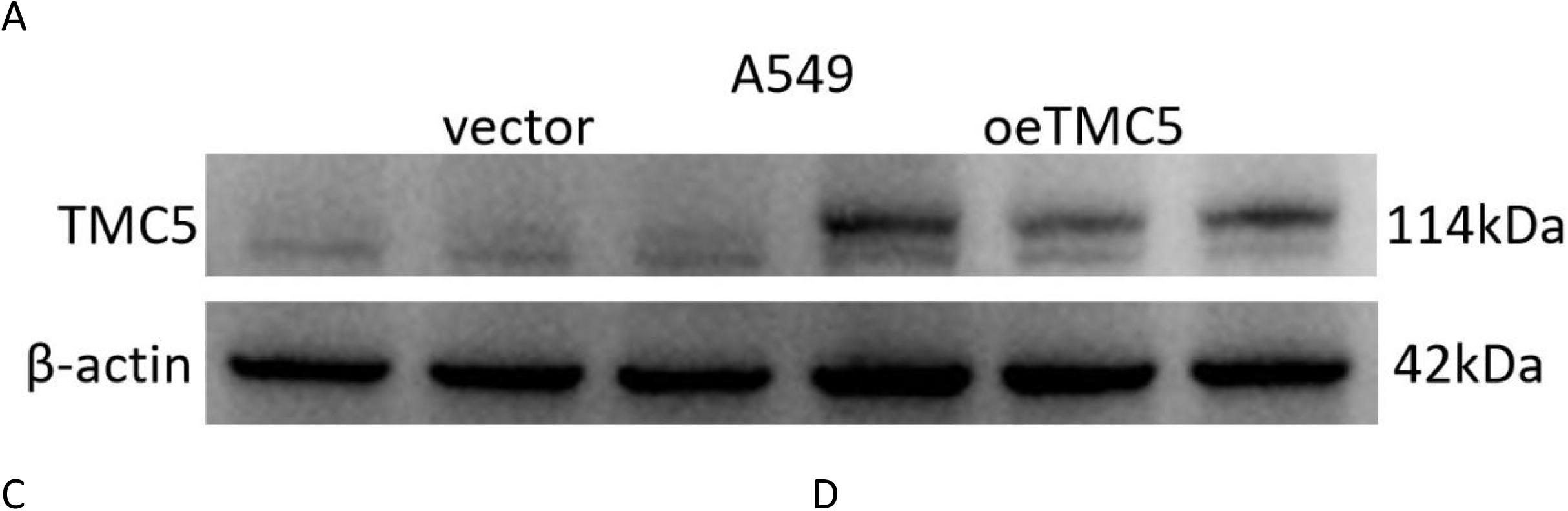
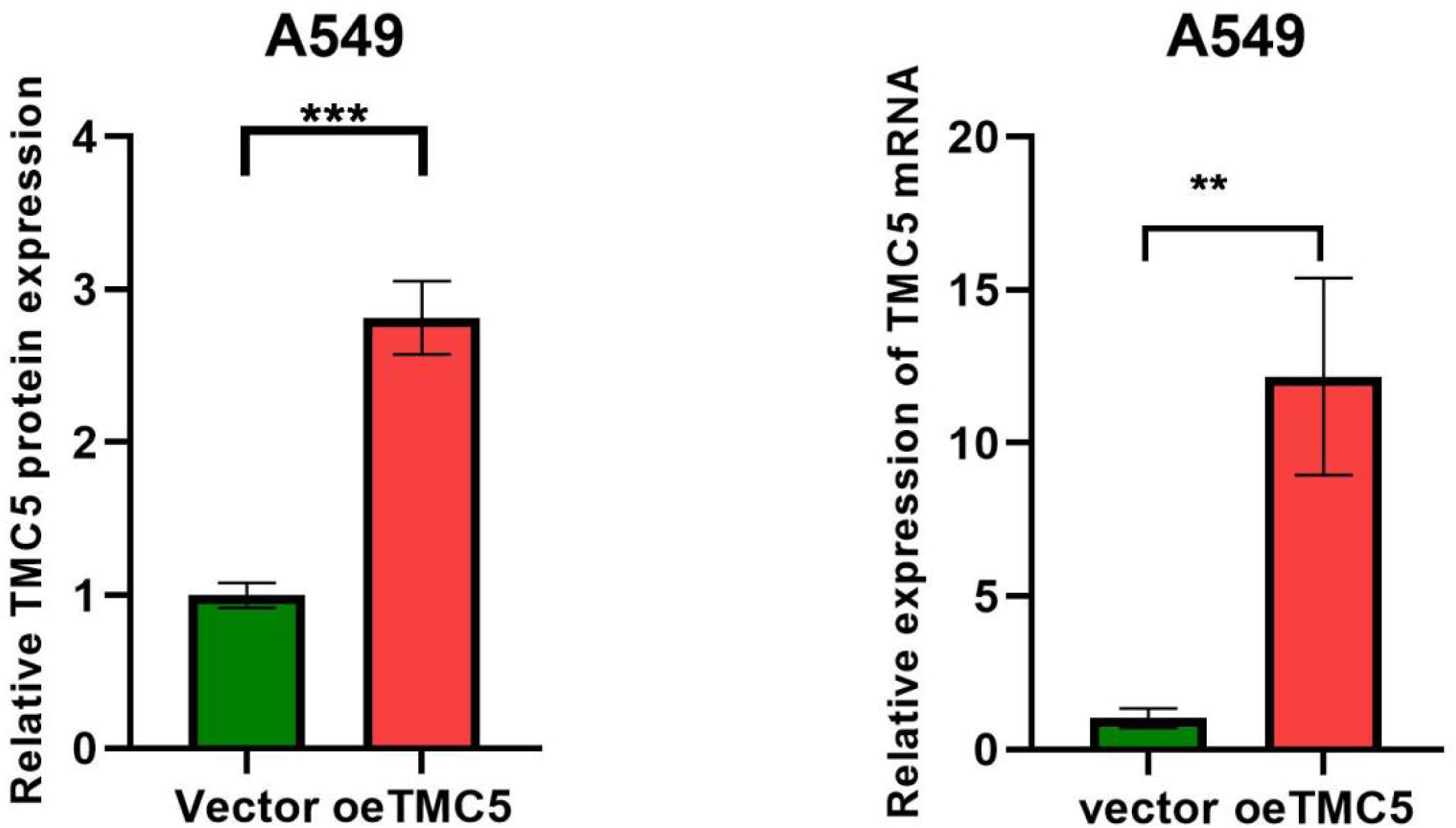
Figure 3. Construction of an A549 cell line overexpressing TMC5.(A,B) Protein level of TMC5 was elevated in the TMC5-overexpressing A549 cells. (C) mRNA level of TMC5 was increased in the TMC5-overexpressing A549 cells. Vector: Empty vector control for overexpression. oeTMC5: Overexpression of TMC5. *P < 0.05, **P < 0.01, ***P < 0.0001.

### TMC5 could promote proliferation and migration of LUAD cells by regulating EMT

To determine whether TMC5 contributes to LUAD cell proliferation and migration by regulating epithelial-mesenchymal transition (EMT), we analysed the protein levels of epithelial (E-cadherin) and mesenchymal (N-cadherin and Vimentin) markers in TMC5-overexpressing A549 and TMC5-knockdown NCI-H1437 cells. We successfully generated stable TMC5-overexpressing A549 cell lines using lentiviral transduction and subsequent puromycin selection(Fig. 4A-C).

TMC5 overexpression down-regulated the expression of E-cadherin and up-regulated the expression of N-cadherin and vimentin in A549 cells (Fig. 5). In contrast, knockdown of TMC5 was associated with strong inhibition of N-cadherin expression and up-regulation of E-cadherin expression in NCI-H1437 cells (Fig. 5). No vimentin protein was detected in NCI-H1437 cells by western blotting. Knockdown of TMC5 was associated with strong inhibition of vimentin expression in A549 cells (Fig. 5). These data suggested that TMC5 regulated proliferation and migration by inducing the EMT program.

**FIGURE 5.**
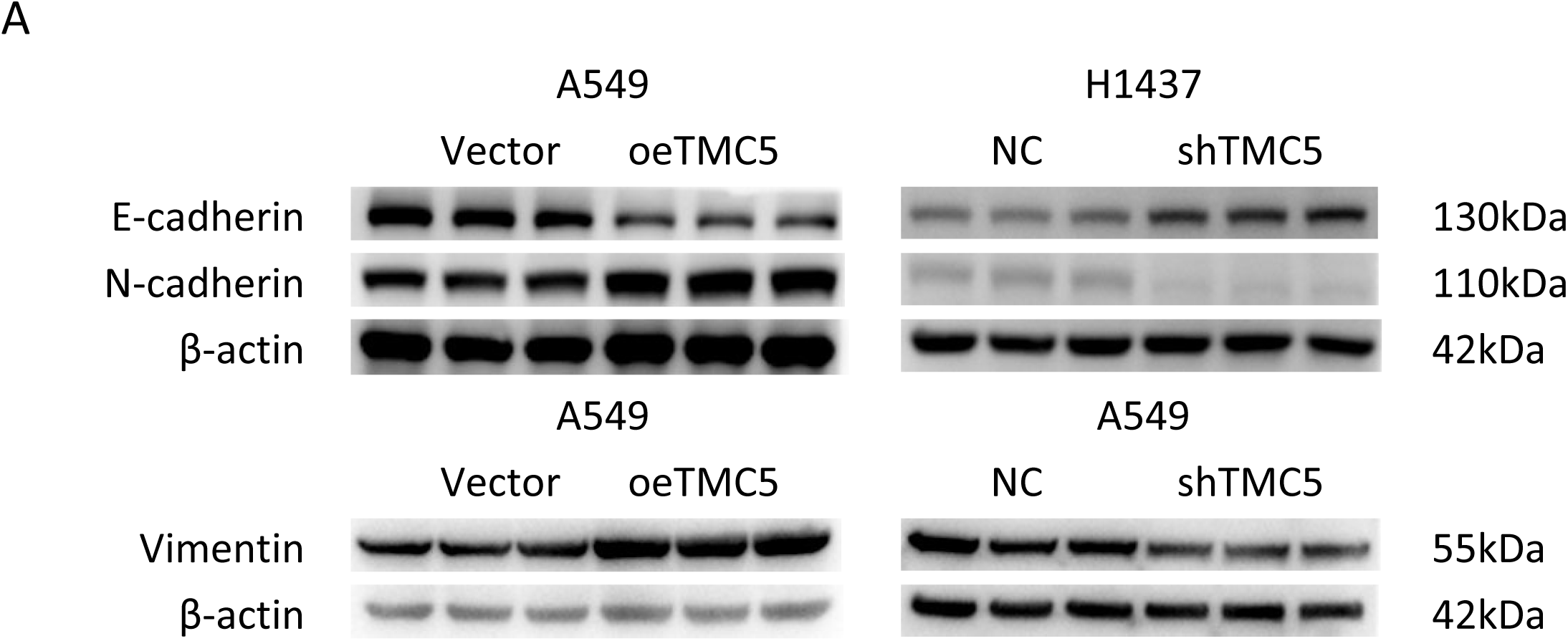
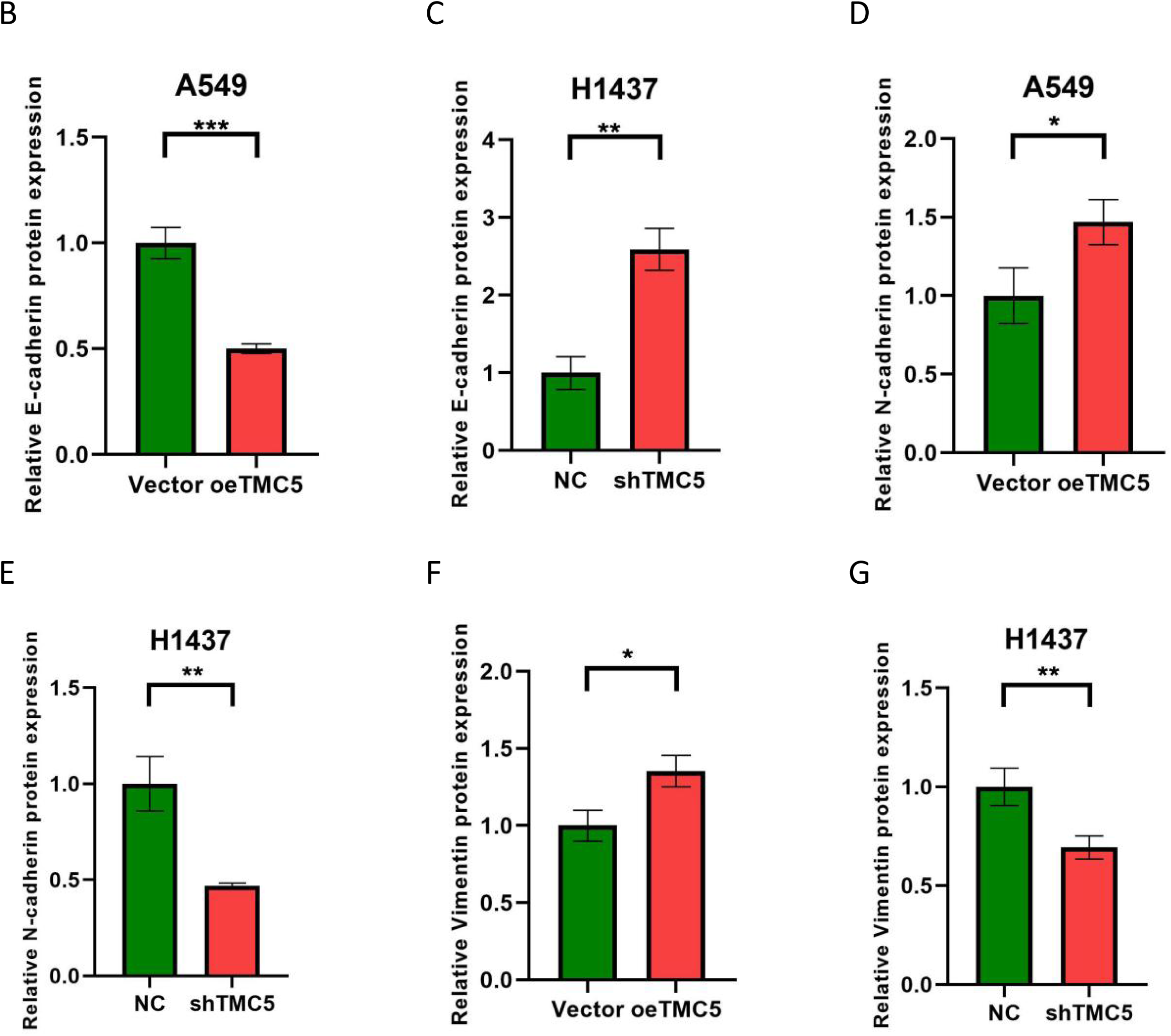
Protein expression of EMT-related markers. In TMC5-knockdown cells, proteins associated with epithelial markers (E-cadherin) were up-regulated, while those related to mesenchymal markers (N-cadherin, Vimentin) were down-regulated. In TMC5-overexpressing cells, proteins linked to epithelial markers (E-cadherin) were down-regulated, whereas those associated with mesenchymal markers (N-cadherin, Vimentin) were up-regulated. EMT, epithelial-mesenchymal transition. (G-L) Statistical analysis showed that the differences were statistically significant. Vector: Empty vector control for overexpression. oeTMC5: Overexpression of TMC5. NC: Normal control. shTMC5: Knockdown of TMC5 via short hairpin RNA (shRNA). *P < 0.05, **P < 0.01, ***P < 0.0001.

## Discussion

The transmembrane channel-like (TMC) family is an evolutionarily conserved group of genes that encode ion channel-like membrane proteins and comprises eight members. TMC5 may play an essential role in the pathogenesis and progression of several diseases. In terms of benign diseases, whole blood transcriptome analysis suggested that TMC5 may be associated with metabolic unhealthy overweight/obesity (7). Numerous bioinformatics analyses have revealed a close association between TMC5 and cancer. Single-cell RNA sequencing (scRNA-seq) data revealed that TMC5 was evidently up-regulated in cancerous epithelial cells of pancreatic adenocarcinoma, and the major predicted TMC5 transcription regulator was STAT3 (8). Proteomic and phosphoproteomic analyses conducted by Zhu et al. (9) revealed that TMC5 was one of the significantly dysregulated differential proteins in colorectal cancer. According to KEGG and GO analysis, TMC5 was involved in the immune response of renal clear cell carcinoma (10). To our knowledge, Zhang et al. (3) first explored the biological functions of TMC5 in cancer through molecular biology experiments. Bioinformatics analysis showed that the mRNA expression level of TMC5 was significantly upregulated in prostate cancer tissues. siRNA-mediated knockdown of TMC5 markedly inhibited the proliferation of prostate cancer cells. Recently, numerous studies have demonstrated that TMC5 plays a significant role in the development and progression of various cancers. The study conducted by Wei et al. (5) suggested the potential of TMC5 as a therapeutic target. In vitro, functional assays demonstrated that TMC5 promoted gastric cancer cell proliferation, migration, and invasion. Animal studies demonstrated that TMC5 promoted gastric cancer growth and metastasis. Cell-based and animal studies have demonstrated that TMC5 facilitated malignant behaviors in colon adenocarcinoma. Additionally, the study conducted by Tian et al. (11) provided the first molecular biological evidence confirming TMC5 as a downstream target of RBM15. TMC5 is highly expressed in hepatocellular carcinoma, and its mechanism of promoting hepatocellular carcinoma progression involves the regulation of epithelial-mesenchymal transition (EMT) (4).

Through analysis of high-throughput data from The Cancer Genome Atlas (TCGA) database, Zhan et al. (12) identified TMC5 as a candidate marker for distinguishing LUAD from lung squamous cell carcinoma. Immunohistochemical results confirmed the potential of TMC5 in differentiating between LUAD and lung squamous cell carcinoma.The expression and function of TMC5 in lung cancer have received little attention and remain unclear. In our study, we performed RT-qPCR on the residual RNA from transcriptome sequencing. Compared with normal lung tissues, TMC5 was significantly overexpressed in lung adenocarcinoma tissues, suggesting its potential as a biomarker for LUAD. We knocked down TMC5 expression in A549 and NCI-H1437 cells using shRNA. Both CCK-8 and colony formation assays showed that cell proliferation was significantly reduced in TMC5-knockdown A549 and NCI-H1437 cells. Transwell assays demonstrated that knockdown of TMC5 led to decreased migration and invasion abilities of A549 and NCI-H1437 cells. These results indicate that TMC5 is an oncogene that promotes the progression of LUAD. Our findings that TMC5 drives proliferation, migration, and invasion in LUAD align with its reported roles in other cancers (3)(5).

Epithelial–mesenchymal transition (EMT) is a process wherein epithelial cancer cells lose their inherent adhesion and transform into more invasive mesenchymal-like cells. As a powerful engine driving tumor metastasis, EMT is the first domino in the multi-step invasion–metastasis cascade of cancer cells (13). Among patients with non-small cell lung cancer, EMT promote cancer development and poor prognosis. During the process of EMT, epithelial markers are downregulated while mesenchymal markers are upregulated. We established A549 cells with stable TMC5 overexpression and NCI-H1437 cells with TMC5 knockdown. In TMC5-overexpressing cells, the epithelial marker E-cadherin was significantly downregulated, whereas in TMC5-knockdown cells, E-cadherin was significantly upregulated. In TMC5-overexpressing cells, mesenchymal markers such as N-cadherin and vimentin were significantly upregulated, while in TMC5-knockdown cells, these mesenchymal markers were significantly downregulated. These results indicate that TMC5 promotes the progression of LUAD by regulating EMT. We establish that, akin to its role in hepatocellular carcinoma as reported by Li et al. (4), TMC5 exerts its pro-tumorigenic effects in LUAD by regulating EMT.

## Conclusion

In summary, our study confirms that TMC5 is highly expressed in LUAD. For the first time, we have demonstrated through cell biology experiments that TMC5 promotes the proliferation, migration, and invasion of LUAD cells. The molecular mechanism by which TMC5 promotes LUAD progression is through the regulation of the EMT process.

## Data availability statement

The data that support the findings of this study are available within the article and its supplementary materials.

## Ethics statement

This study was performed in conformity with the Declaration of Helsinki. It was approved by the Ethics Committee of the First People’s Hospital of Yunnan Province (Authorization number: KHLL2024-KY282).

## Author contributions

Danxiong Sun: Writing – original draft. Rufang Li: Writing – original draft. Yunhui Zhang: Writing – review & editing.

## Generative AI statement

The author(s) declare that no Generative AI was used in the creation of this manuscript.

